# A Novel Water-Soluble Photosensitizer for Photodynamic Inactivation of Gram-Positive Bacteria

**DOI:** 10.1101/2020.05.29.124768

**Authors:** Zihuayuan Yang, Ying Qiao, Junying Li, Fu-Gen Wu, Fengming Lin

## Abstract

Antimicrobial photodynamic therapy (APDT) is a promising alternative to traditional antibiotics for bacterial infections, which inactivates a broad spectrum of bacteria. However, it has some disadvantages including poor water solubility and easy aggregation of hydrophobic photosensitizers (PS), and poor tissue penetration and cytotoxicity when using UV as the light source, leading to undesired photodynamic therapy efficacy. Herein, we develop a novel water-soluble natural PS (sorbicillinoids) obtained by microbial fermentation using recombinant filamentous fungus *Trichoderma reesei* (*T*. *reesei*). Sorbicillinoids could effectively generate singlet oxygen (^1^O_2_) under ultraviolet (UV) light irradiation, and ultimately display photoinactivation activity on Gram-positive bacteria, but not Gram-negative ones. *Staphylococcus aureus* (*S*. *aureus*) treated with sorbicillinoids and UV light displayed high levels of intracellular reactive oxygen species (ROS), notable DNA photocleavage, and compromised cell semi-permeability without overt cell membrane disruption. Moreover, the dark toxicity, phototoxicity or hemolysis activity of sorbicillinoids is negligible, showing its excellent biocompatibility. Therefore, sorbicillinoids, a type of secondary metabolite from fungus, has a promising future as a new PS for APDT using nontoxic dose of UV light irradiation.

**Importance:** It is of great value to develop novel PSs for APDT to enhance its efficacy for the reason that many traditional PSs have disadvantages like low water solubility and poor biocompatibility. In this study, we develop a novel water-soluble natural PS - sorbicillinoids obtained by microbial fermentation using T. reesei. Sorbicillinoids could effectively generate singlet oxygen under UV light irradiation, and ultimately display photoinactivation activity on Gram-positive bacteria, but not Gram-negative ones. More importantly, UV light can generally only be used to inactivate bacteria on the surface due to its weak penetration. However, it can penetrate deep into the solution and inactivate bacteria in the presence of sorbicillinoids. Therefore, sorbicillinoids, a type of secondary metabolite from fungus, has a promising future as a new PS for APDT using nontoxic dose of UV light irradiation.

## Introduction

Diseases caused by bacterial infections, such as pneumonia and sepsis, threaten the lives of millions of people every year (1,2). The most widely used strategy for coping with bacterial infections at present is antibiotics (3). However, the abuse of antibiotics in recent decades has led to an increase in the resistance of bacteria (4), which causes the emergence of multi-drug resistant (MDR) bacteria (5,6). Thus, it is urgent to develop new antibiotics and efficient antibacterial therapies to fight MDR bacteria. Many novel antibiotics have been explored to successfully combat MDR bacterial infection, including carbon dots nanoparticles (NP) (7,8) and cationic antibacterial peptides (9,10). In addition, new efficient antibacterial therapies have been also applied in treating bacterial infections, such as photodynamic therapy (PDT) (11,12) and photothermal therapy (PTT) (13,14). Antimicrobial photodynamic therapy (APDT) has been widely reported as a promising alternative to traditional antibiotics in the past few years (15,16). APDT generally involves molecular oxygen, light source, and photosensitizer (PS). With light irradiation of a suitable wavelength, PS converts molecular oxygen into reactive oxygen species (ROS) (12), primarily singlet oxygen (^1^O_2_). The highly active ROS can cause damage to important biomolecules such as lipids (17), proteins (18), and DNA (19), ultimately leading to bacterial cell death. Unlike conventional antibiotics, APDT only irradiates the lesion location and does not cause damage to other parts (11), thereby achieving selective bacterium-killing. More importantly, APDT can inactivate a broad spectrum of bacteria, opening new avenues for the development of new antibacterial therapies (12).

Most currently available PSs for APDT have poor solubility in water, showing a strong tendency to aggregate, ultimately leading to poor photodynamic therapy efficacy (20). To improve their hydrophilicity, a variety of modification methods were applied by physical encapsulation or chemical covalent conjugation of PSs to nanocarriers such as peptide nanoparticles (21), polymer nanoparticles (22), and liposomes (23). However, these surface modification methods are time-consuming and usually bring some unexpected consequences such as low biocompatibility, which may compromise the clinical application of APDT. Another issue that plagues APDT was the need of UV light for some PSs like metal oxide NPs zinc oxide (ZnO) NPs and titanium dioxide (TiO_2_) NPs (24), for the reason that UV light is cytotoxic to human cells and has poor tissue penetration (25). Therefore, it is urgent to find new PSs for the APDT system to overcome these issues. Natural products from fungus and plants have been reported as a promising alternative to conventional PSs such as hypericin (26), curcumin (27), hypocrellin (28) and cationic riboflavin (29). The compounds are usually derived from secondary metabolism. Natural products as PSs have the advantages of good biocompatibility, wide source, and high ^1^O_2_ yield (30).

Sorbicillinoids are hexaketide metabolites isolated from both marine and terrestrial ascomycetes, such as *Trichoderma*, *Acremonium*, *Aspergillus*, *Emericella*, *Penicillium*, *Phaeoacremonium*, and *Scytalidium* (31). Most of these compounds have characteristic structures, including bicyclic or tricyclic structures and C1’–C6’ sorbyl sidechain. Sorbicillinoids were divided into four classes based on their structure (32): monomeric sorbicillinoids, bisorbicillinoids, trisorbicillinoids, and hybrid sorbicillinoids. In recent years, various potential applications of sorbicillinoids have been exploited, such as antioxidants (33,34), antibiotics (35,36), and anticancer drugs (37,38). However, research on sorbicillinoids as a PS for the APDT system has not been reported.

In this study, we discovered a new type of natural material as a water-soluble PS for APDT. Sorbicillinoids produced by *Trichoderma reesei* converts oxygen molecules into ^1^O_2_ effectively under mild ultraviolet (UV) irradiation. As a result, sorbicillinoids exhibited great APDT effect toward Gram-positive bacteria. The underlying mechanism of the light-activated antibacterial activity of sorbicillinoids was explored by monitoring intracellular ROS, DNA photocleavage and cell membrane damage. Furthermore, the biocompatibility of sorbicillinoids-mediated APDT was evaluated using MTT assay and hemolysis assay.

## Results and Discussions

### Singlet oxygen generation

Sorbicillinoids was produced from the fermentation of the recombinant *Trichoderma reesei* (*T. reesei*) strain ZC121 in *Trichoderma* minimal media (39) (TMM) with an excellent sorbicillinoids production ability. The obtained sorbicillinoids mainly included sorbicillinol, bisvertinolone and oxosorbicillinol as identified by LC-MS (40). Given that sorbicillinoids has great absorbance at 370 nm (41), there is a possibility that sorbicillinoids can generate ^1^O_2_ under light irradiation. To prove this, the ^1^O_2_ generation ability of sorbicillinoids under the irradiation of both UV and white light was measured using a SOSG kit (Figure 1). Sorbicillinoids produced significant ^1^O_2_ when irradiated with UV light. The produced ^1^O_2_ was increased noticeably with the irradiation time increasing, being 300% that of UV alone at 30 min. In contrast, when sorbicillinoids was exposed to white light for 30 min, insignificant ^1^O_2_ production was observed. No ^1^O_2_ generation was observed for sorbicillinoids alone. This result confirmed our hypothesis that sorbicillinoids can synthesize ^1^O_2_ in the presence of UV light, but not visible light. It is worth noting that ^1^O_2_ is highly toxic to the microorganisms, serving as one major factor for APDT’s antibacterial effect (15,42). Therefore, sorbicillinoids from *T. reesei* might be explored as a new PS for APDT to kill microorganisms.

**Figure 1.**
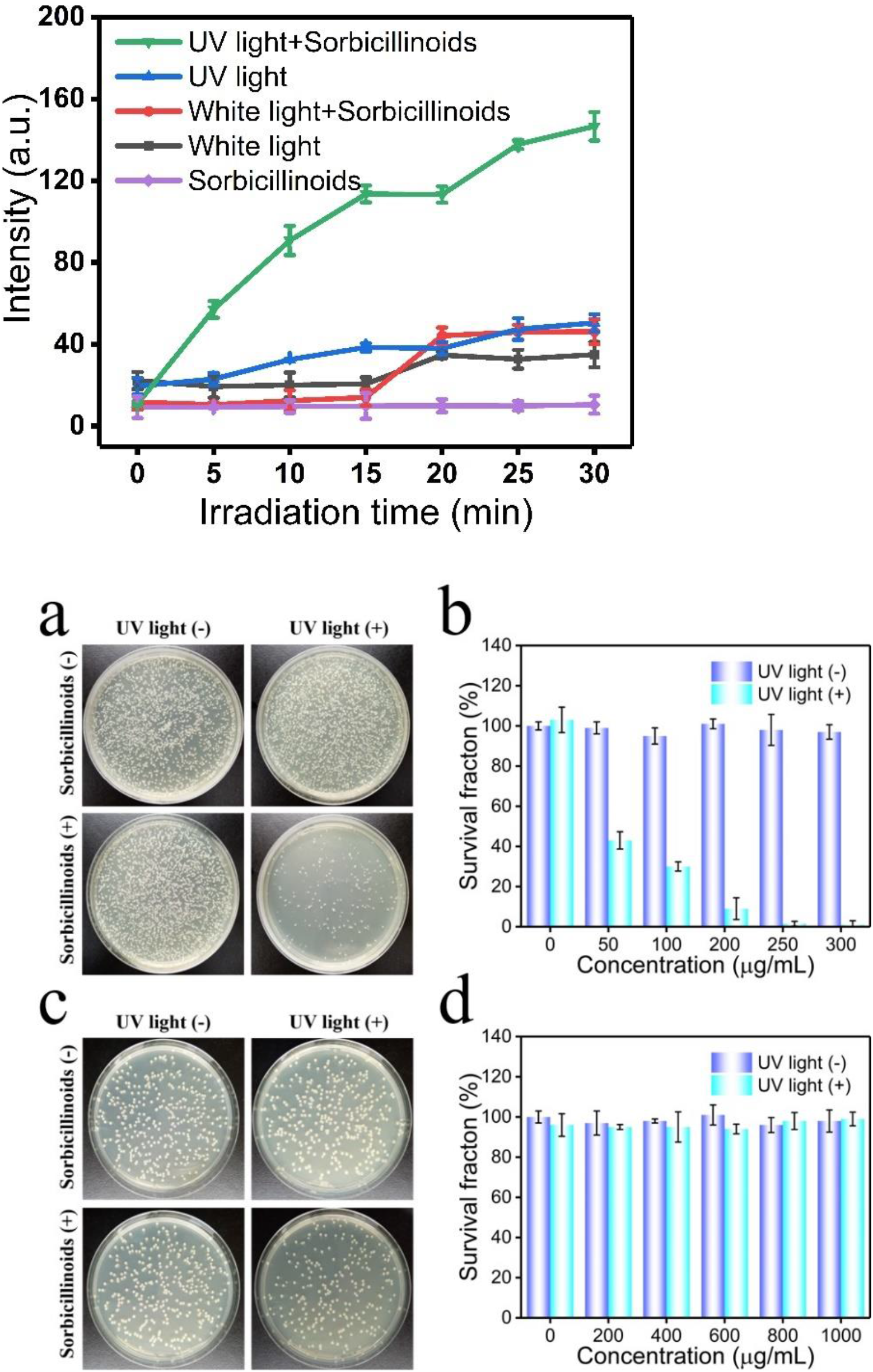
^1^O_2_ generation of sorbicillinoids in 0.9% NaCl as measured by the fluorescence intensity changes of SOSG at 528 nm with the excitation wavelength of 504 nm.

### Antibacterial activity

Inspired by their excellent ^1^O_2_ production ability under UV irradiation, we investigated the APDT ability of sorbicillinoids. *S*. *aureus* and *E*. *coli* representing Gram-positive and Gram-negative bacteria respectively, were incubated with different concentrations of sorbicillinoids in 1.5 mL sterilized centrifuge tubes, followed by UV light irradiation for 30 min (Figure 2). When irradiated under UV light, *S*. *aureus* was killed by 60.3%, 68.5%, 90.5%, 98.1%, and 99.5% in the presence of 50, 100, 200, 250, and 300 μg/mL sorbicillinoids, respectively (Figure 2a and 2b). Conversely, no antibacterial effect of sorbicillinoids was observed on *E. coli*, even when the administrated concentration was increased up to 1000 μg/mL (Figure 2c and 2d). Sorbicillinoids or UV alone was non-toxic to both *S*. *aureus* and *E. coli* (Figure 2). The undetected antibacterial effect of UV light in this study is probably ascribed to the poor penetration ability of V light through the plastic centrifuge tube (43). It appears that while the plastic is mitigating the intensity of incident UV light to the point where there is no cell death in the negative control, there is still enough incident UV light to sensitize sorbicillinoids to generate drastic ^1^O_2_, leading to the killing of *S*. *aureus*.

**Figure 2.**
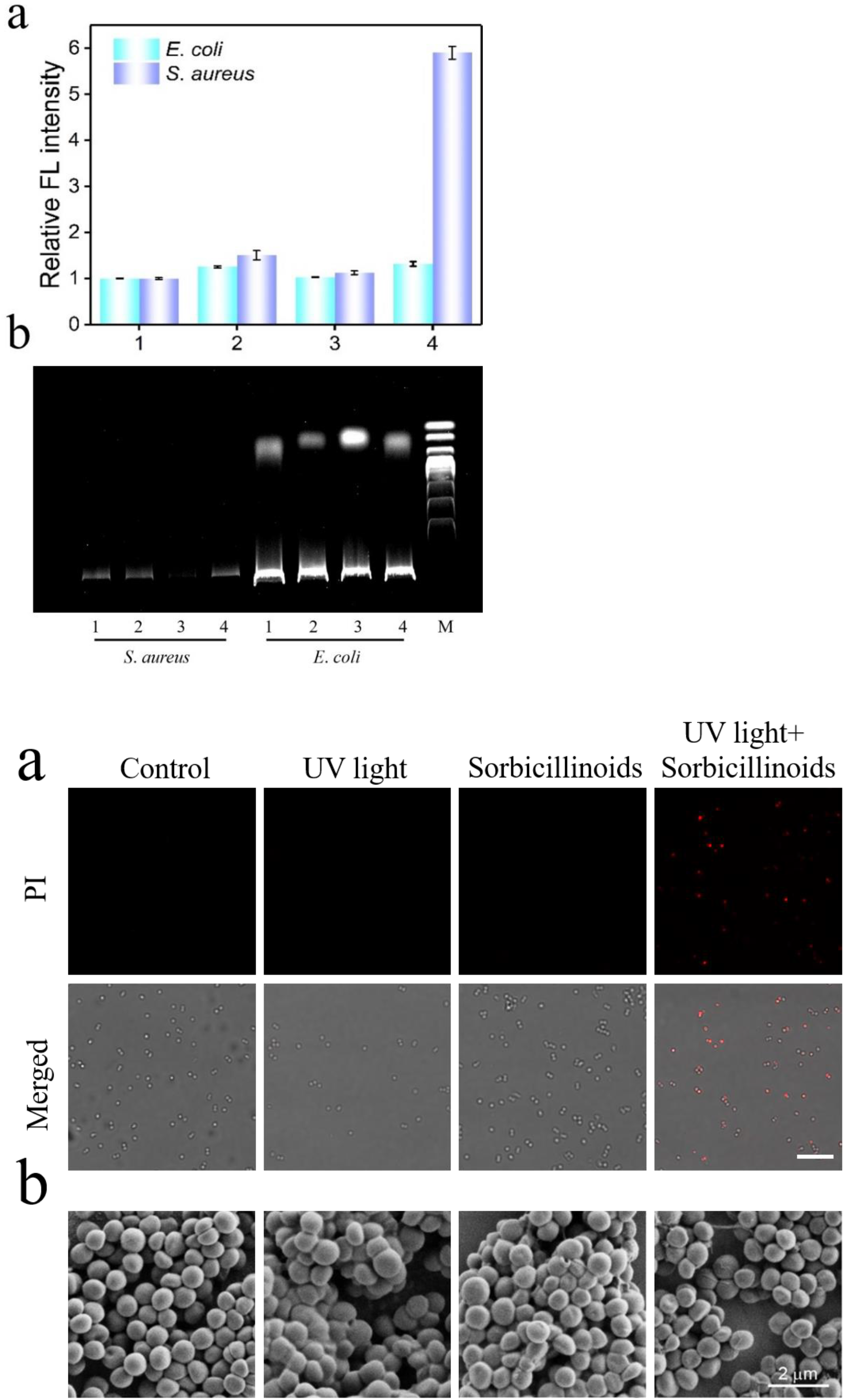
The APDT ability of sorbicillinoids against *S*. *aureus* (a, b) and *E*. *coli* (c, d). Agar plate photographs of *S*. *aureus* (a) and *E*. *coli* (c) treated with sorbicillinoids under (no) UV light irradiation conditions. The corresponding dependence of bacterial survival fraction on the concentration of sorbicillinoids was measured (b, d).

Moreover, the UV-mediated APDT ability of sorbicillinoids was tested on another two Gram-positive bacteria *B*. *subtilis* and *M*. *luteus*, and another Gram-negative bacterium *P*. *vulgaris*. In the presence of 25, 50, 75, 100, 200 and 300 μg/mL sorbicillinoids and under UV light irradiation, *B*. *subtilis* was killed by 36.7%, 40.4%, 61.4%, 89.3% and 98.5% and 98.9%, respectively, whereas *M*. *luteus* was killed by 29.5%, 80.8%, 98.0%, 99.7% and 99.9%, respectively in the presence of 50, 100, 200, 250 and 300 μg/mL sorbicillinoids (Figure S1). By contrast, no antibacterial ability was observed against *P*. *vulgaris* (Figure S2) as it is in the case of *E*. *coli*. Obviously, sorbicillinoids with the irradiation of nontoxic UV light can successfully kill Gram-positive bacteria, but not Gram-negative ones. This Gram-selective antibacterial ability of sorbicillinoids may be attributed to the different cell wall structures of Gram-positive bacteria and Gram-negative bacteria. It is well-known that compared to Gram-positive bacteria, Gram-negative bacteria have an outer membrane that lies outside of the plasma membrane, serving as a penetration barrier and protecting Gram-negative bacteria from antibiotics and PSs (10,44). This outer membrane may prevent sorbicillinoids from entering Gram-negative cells to realize APDT. The localization of PSs largely determines the efficiency of PDT, as the half-life of ^1^O_2_ is very short (< 0.04 μs) and its action radius is only shorter than 0.02 μm in the biological system (45). Therefore, despite it could produce abundant ^1^O_2_ in the solution under UV light irradiation, the extracellular sorbicillinoids exhibited no bactericidal activity against Gram-negative bacteria. For Gram-positive bacteria without the outer membrane, the water-soluble sorbicillinoids with small molecular weight might readily diffuse through cell surface into the cytosol to exert APDT.

### Intracellular ROS generation and agarose gel electrophoresis of genomic DNA

To evidence our speculation that sorbicillinoids enters Gram-positive bacteria to realize APDT, the generation of intracellular ROS was tested employing 2’,7’-dichlorodihydrofluorescein diacetate (DCFH-DA) probe. DCFH-DA freely passes through the cell membrane into cells where it is hydrolyzed by intracellular esterase to form 2’,7’-dichlorodihydrofluorescein (DCFH) that cannot penetrate the cell membrane and stay inside cells. DCFH is subsequently oxidized by the intracellular ROS into 2’,7’-dichlorofluorescein (DCF) that has fluorescence. Neither DCFH-DA nor DCFH has fluorescence. Therefore, the level of ROS in the cells can be measured by detecting the fluorescence of DCF. After treated with sorbicillinoids and irradiated under UV light, the fluorescence intensity of *S*. *aureus* increased significantly by 5 times as compared to the untreated *S*. *aureus*, whereas no increasing fluorescence intensity was observed for *E*. *coli* (Figure 3a). This suggested that the treatment of sorbicillinoids with the irradiation of UV induced *S*. *aureus* to generate large sums of intracellular ROS, which might further damage biomacromolecules like proteins, DNA and lipids. This result matched well with the selective photoinactivation of sorbicillinoids against Gram-positive bacteria as we observed above.

**Figure 3.**
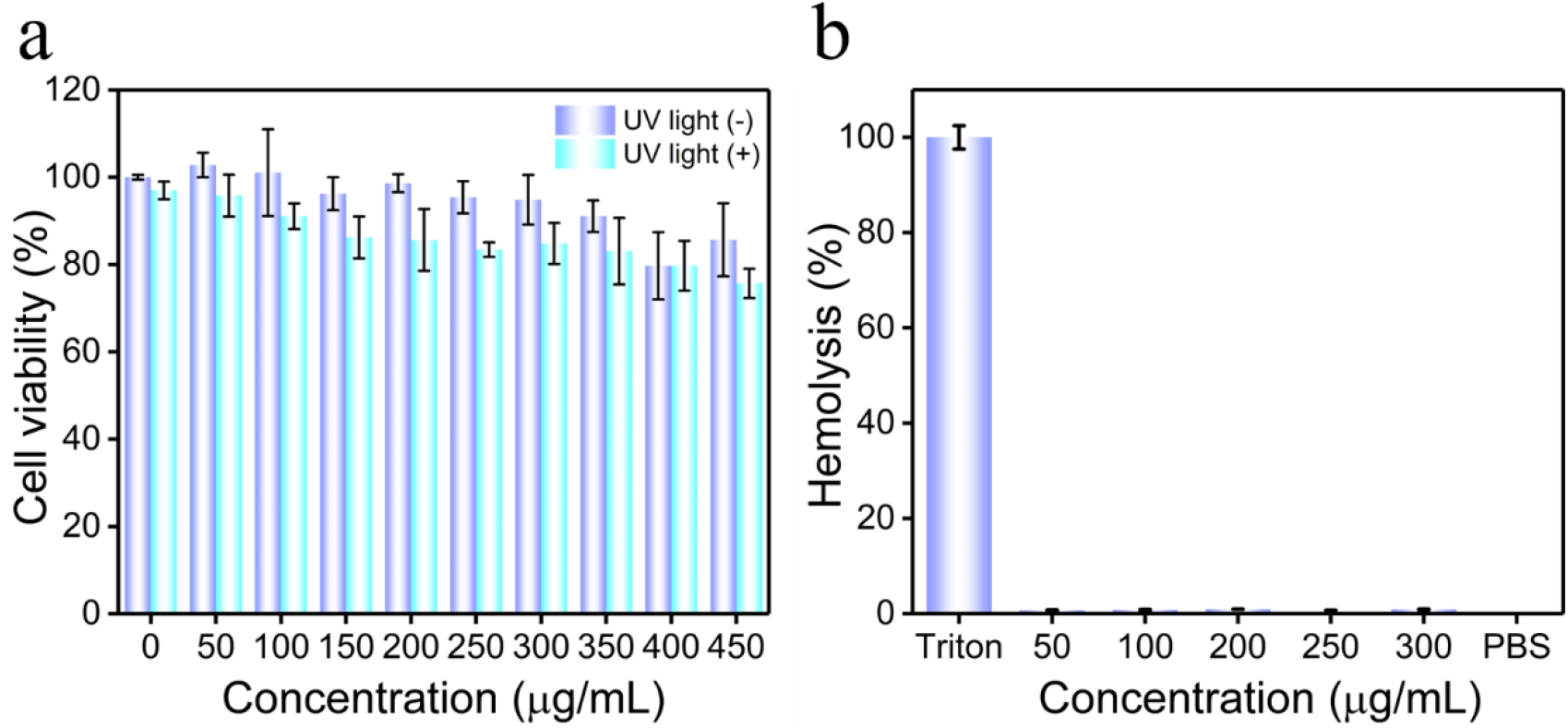
a) Generation of intracellular ROS in *S*. *aureus* and *E*. *coli*. b) Electrophoretic profiles of DNA extracted from *S*. *aureus* and *E*. *coli*. 1: no treatment; 2: UV light; 3: sorbicillinoids; 4: sorbicillinoids and UV light; M: molecular weight marker.

Inspired by its ultra-high intracellular ROS level, DNA photocleavage of *S*. *aureus* with the treatment of sorbicillinoids and UV irradiation was detected by agarose gel electrophoresis (Figure 3b). There was a marked reduction of DNA for *S*. *aureus* treated with 100 μg/mL sorbicillinoids under UV light irradiation, indicating nucleic acid photocleavage took place in the treated bacterial cells. *S*. *aureus* treated with sorbicillinoids or UV light alone did not display a reduction of DNA. On the contrary, sorbicillinoids did not result in observable DNA reduction in *E*. *coli* with or without UV light. Obviously, the unusually high intracellular ROS induced by sorbicillinoids upon UV irradiation caused cellular DNA damage in *S. aureus*.

### PI staining and morphological characterization of bacteria

Both DNA and cytoplasmic membrane are two crucial cellular components targeted by ROS in APDT (46). Therefore, the effect of sorbicillinoids on the cell membrane of *S*. *aureus* and *E*. *coli* was investigated using both propidium iodide (PI) staining and SEM imaging (Figure 4 and Figure S3). PI cannot penetrate into viable bacteria cells with complete membrane. Yet it can enter bacterial cells with damaged membrane to intercalate between the base pairs of the accessible double-stranded DNA and emit red fluorescence with the excitation at 580 nm. *S*. *aureus* were dyed red by PI after the treatment of sorbicillinoids and UV irradiation (Figure 4a), demonstrating their membrane permeability was compromised. Nevertheless, no red fluorescence was observed in the irradiated *E*. *coli* cells in the presence of sorbicillinoids, showing their cell membrane was not affected (Figure S3a). This finding was consistent with that sorbicillinoids exhibited no APDT activity against *E. coli*. Similar to most classical PSs (46), sorbicillinoids induces oxidative damage to the cell membrane of *S*. *aureus*, contributing to its overall antimicrobial activity together with the high intracellular ROS level and the notable DNA photocleavage.

**Fig 4.**
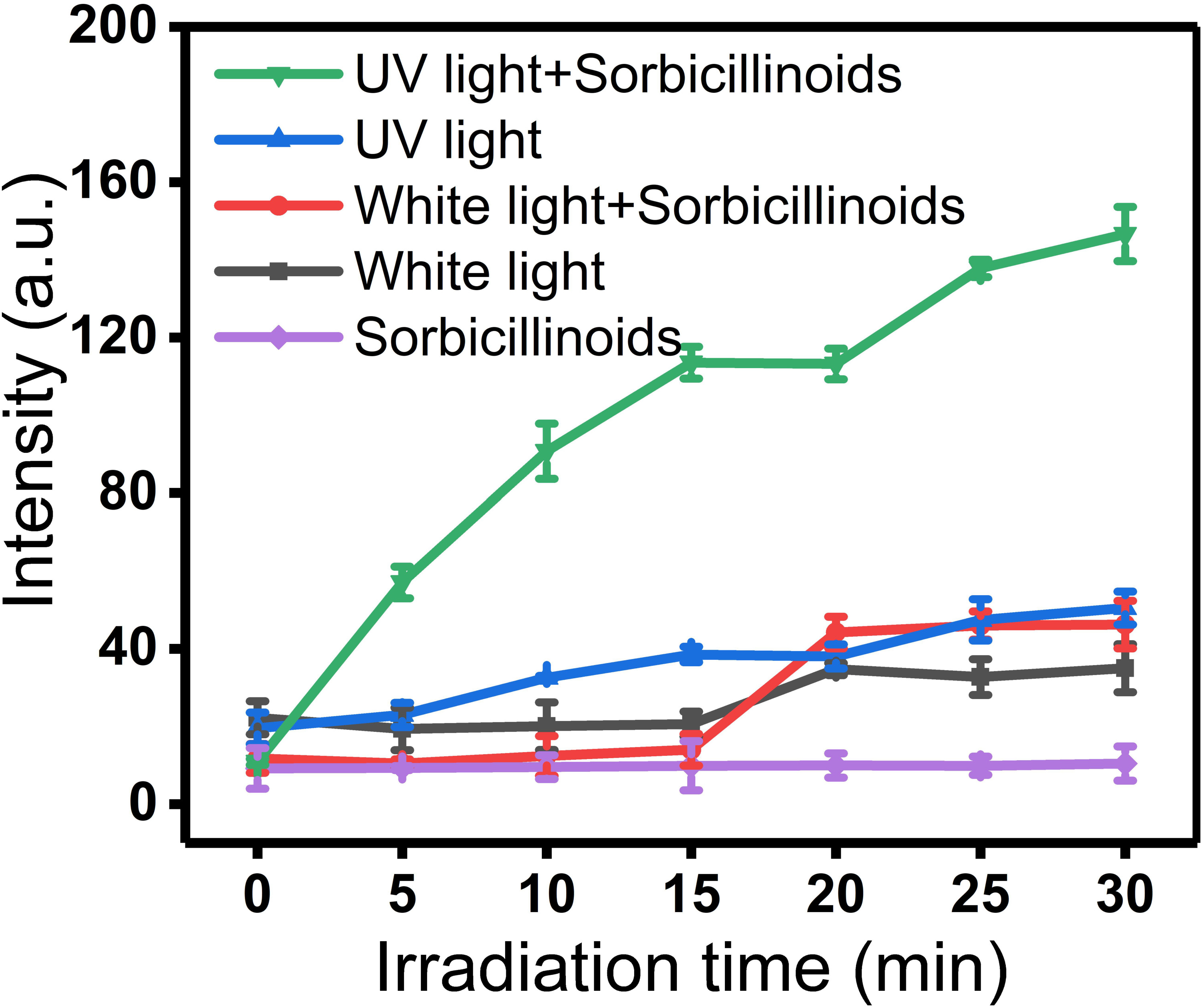
a) Confocal fluorescence images of *S*. *aureus* that was incubated with PI after different treatments as indicated in the figure. Bacteria with damaged cell membrane would be stained by PI, showing red fluorescence. (scale bar = 10 μm). b) the corresponding SEM images were also taken.

Furthermore, SEM experiments were carried out to monitor the impact of sorbicillinoids on the morphology of bacterial surface. No noticeable morphology change was found in the SEM pictures of the irradiated *S. aureus* (Figure 4b) and *E. coli* (Figure S3b), including wrinkles, holes or leaked intracellular substances as reported previously for some photodynamic antibacterial materials (15,47). Sorbicillinoids killed *S. aureus* without overt cell membrane disruption, as in the case of other PSs like EPS-RB NPs (48). Taken together, sorbicillinoids can readily diffuse through the cell wall of Gram-positive bacteria into the cells. When irradiated under UV light, it produced large sums of intracellular ROS, thereby targeting and destroying both cytoplasmic membrane and DNA to achieve APDT effect. This does not happen to Gram-negative bacteria due to their distinctive outer membrane that is an extra effective penetration barrier to block sorbicillinoids out of cells.

### Cytotoxicity evaluation assay and hemolysis activity

Last but not least, the biocompatibility of sorbicillinoids was assessed by the cytotoxicity measurement and hemolysis assay (Figure 5). Sorbicillinoids showed a little cytotoxicity to AT-II cells at 400 μg/mL (Figure 5a) that was higher than its bactericidal concentrations for *S*. *aureus* (Figure 4a), *B*. *subtilis* (Figure S1a) and *M*. *luteus* (Figure S1b). When the concentration of sorbicillinoids was increased to 450 μg/mL, more than 85% of AT-II cells remained alive, demonstrating a high safety of sorbicillinoids (Figure 5a). More importantly, even under the irradiation of UV light, only a slight decrease of the cell viability was found with > 75.66% AT-II cells alive when the concentration of sorbicillinoids was increased up to 450 μg/mL. Thus, sorbicillinoids did not induce marked dark toxicity or phototoxicity to AT-II cells. On the other hand, only 0.89% hemolysis was observed with the treatment of 300 μg/mL sorbicillinoids (Figure 5b and Figure S4). Sorbicillinoids show excellent biocompatibility, which is of benefit to their future practical applications in clinic.

**Figure 5.**
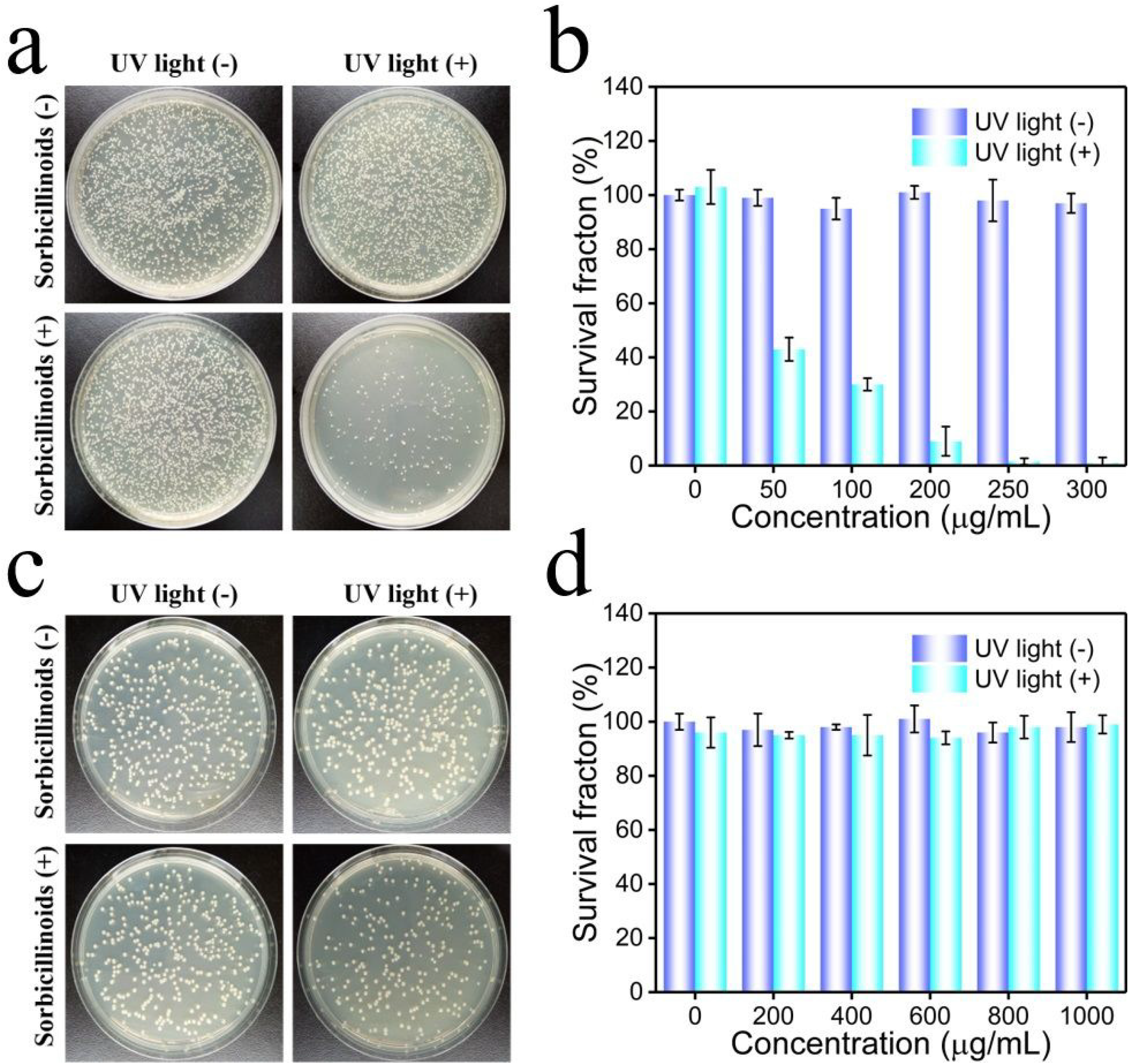
Biocompatibility evaluation of sorbicillinoids. a) Viabilities of AT-II cells in the presence of different concentrations of sorbicillinoids with and without UV light. b) Hemolysis rates of RBCs after incubation with various concentrations of sorbicillinoids. RBCs in Triton X-100 and PBS were set as the positive and negative controls, respectively.

## Materials and Methods

### Materials

Luria Bertani (LB) broth, LB agar and potato dextrose agar (PDA) were purchased from Beijing Land Bridge. Singlet oxygen sensor green (SOSG) kit was acquired from Molecular Probes Inc. (Eugene, Oregon, USA). Propidium iodide (PI) was obtained from KeyGen Biotech (Nanjing, China). Glutaraldehyde, 3-(4,5-dimethyl-2-thiazolyl)-2,5-diphenyl-2-*H*-tetrazolium bromide (MTT), methanol, ethanol, dimethyl sulfoxide (DMSO), and triton X-100 were purchased from Aladdin Chemistry Co., Ltd. (Shanghai, China). Reactive oxygen species assay kit was bought from Beyotime Biotech Inc. (Shanghai, China). All other chemicals used in this study were purchased from Sigma-Aldrich (St. Louis, MO, USA). All solutions were prepared with deionized water (18.2 MΩ cm) purified by a Milli-Q water purification system (Milli-Q, Millipore, USA).

### Sorbicillinoids production

The conidia produced by *Trichoderma reesei* strain ZC121 grown on PDA plates for 7 days at 28 °C, were inoculated into 10 mL sabouraud dextrose broth (SDB) and incubated for 48 h with 200 rpm at 28 °C. Pre-grown mycelia were inoculated with an inoculation ratio of 10% (v/v) into 50 mL TMM with 2% glucose, and then incubated for 120 h with 200 rpm at 28 °C. The suspension was centrifuged at 14000 rpm for 15 min at 4 °C to remove *Trichoderma reesei* cells and other solid materials, and the supernatant was dried at 120 °C in an oven. The obtained powder was dissolved in methanol to remove inorganic salt. Methanol was evaporated at 60 °C under a nitrogen atmosphere to obtain sorbicillinoids. TMM medium was composed of the following chemicals (all concentration unit is g/L unless otherwise noted): (NH_4_)_2_SO_4_, 4.0; KH_2_PO_4_, 6.5; Tween-80, 0.0186% (v/v); Yeast extract, 0.75; Tryptone, 0.25; Maleic acid, 11.6; FeSO_4_·7H_2_O, 0.005; MnSO_4_·H_2_O, 0.0016; ZnSO_4_·7H_2_O, 0.0014; CoCl_2_·6H_2_O, 0.002; MgSO_4_, 0.60; CaCl_2_, 0.60; urea, 1.0. Then the pH of TMM was adjusted to 5.8 by NaOH.

### Singlet oxygen generation

The generation of ^1^O_2_ was measured with singlet oxygen sensor green (SOSG). 1 μL SOSG dissolved in methanol was mixed with 2 mL 0.9% NaCl and 50 μg/mL sorbicillinoids in 0.9% NaCl, followed by UV light (2 mW/cm^2^) or white light irradiation (8 mW/cm^2^). 0.9% NaCl without any light irradiation was set as control. Then, the fluorescence intensity of the samples at 528 nm was measured by a spectrofluorophotometer (RF-5301PC, Shimadzu, Japan).

### Antibacterial activity

*Escherichia coli* (*E*. *coli*), and *Staphylococcus aureus* (*S*. *aureus*) were selected to present Gram-negative and Gram-positive bacteria, respectively. *E*. *coli* or *S*. *aureus* were cultivated overnight in LB medium at 37 °C with 200 rpm. Bacteria were obtained from bacteria suspension overnight by centrifugation at 8000 rpm for 5 min and then resuspended in 0.9% NaCl. The optical density of the resuspension at 600 nm (OD_600_) was measured using a UV-vis spectrophotometer (UV-2600, Shimadzu, Japan). Then, the resuspension was diluted to OD_600_ = 0.5 with 0.9% NaCl. After the dilution, 100 μL bacteria suspension was mixed with 900 μL sorbicillinoids dissolved in 0.9% NaCl at different concentrations. The final concentrations of sorbicillinoids were 0, 50, 100, 200, 250, and 300 μg/mL. for *S*. *aureus* (0, 200, 400, 600, 800, and 1000 μg/mL. for *E*. *coli*). After irradiation under UV light (2 mW/cm^2^) for 30 min, 100 μL solution was plated on LB agar plates. The plates were placed in an incubator at 37 °C for 24 h, followed by colony counting. Antibacterial activity of sorbicillinoids against Gram-negative bacterium *Proteus vulgaris* (*P*. *vulgaris*), and Gram-positive bacteria *Bacillus subtilis* (*B*. *subtilis*) and *Micrococcus luteus* (*M*. *luteus*), were also tested in the same way.

### Intracellular ROS generation

2,7-dichlorofluorescein diacetate (DCFH-DA) was applied to measure the generation of intracellular ROS. 0.5 μL DCFH-DA was added into 500 μL bacteria suspension (OD_600_ =0.05), followed by incubation at 37 °C for 10 min. After centrifugation at 8000 rpm for 5 min, bacteria were resuspended in 500 μL water, supplemented with 5 μL sorbicillinoids (10 mg/mL for *S*. *aureus* and 40 mg/mL for *E*. *coli*) dissolved in water. After incubation at 37 °C for 30 min, the suspension was irradiated under UV light (2 mW/cm^2^) for 30 min. The fluorescence intensity was measured by a flow cytometer (NovoCyte™ 2060, ACEA, USA). Channel used for analyses was FITC with the excitation at 488 nm.

### Agarose gel electrophoresis of bacterial genomic DNA

The genomic DNA of bacteria was extracted with the Bacterial DNA kit which was bought from TIAGEN Biotech Co., Ltd. (Beijing, China). The bacteria cell (1 × 10^6^ CFU/mL) with different treatments as indicated in the antibacterial activity assay were collected by centrifugation at 8000 rpm for 5 min and then washed twice with 0.9% NaCl. The genomic DNA was extracted according to manufacturer’s instruction. Briefly, the bacterial cells were separately resuspended in 200 μL GA buffer, 20 μL proteinase K solution and 220 μL GB buffer. After incubation at 70 °C for 10 min, 220 μL ethanol was added into each sample. Then the solution was transferred to a spin column CB3 and washed with GD buffer and PW washing solution. After that, the genomic DNA was bound onto the spin column and then eluted with TE buffer. The extracted DNA was analyzed by electrophoresis in Tris Acetate-EDTA (TAE) buffer containing 0.8% agarose (w/v) for 50 min at 100 V. The electrophoretic profiles were acquired with a Gel imaging system (Tanon 3500R, Shanghai, China).

### PI Staining

The bacterial cells (1 × 10^6^ CFU/mL) with different treatments as indicated in the antibacterial activity assay were incubated at 37 °C for 2 h and then collected by centrifugation at 8000 rpm for 5 min. After the centrifugation, the bacteria were stained with red-fluorescent nucleic acid stain (PI) for 30 min. The bacteria samples were imaged using a confocal microscope (TCS SP8, Leica, Germany).

### Morphological Characterization of Bacteria

Morphological characterization of bacteria was carried out with a scanning electron microscope (SEM, ULTRA Plus, Zeiss, Germany). The bacterial cells (1 × 10^6^ CFU/mL) with different treatments as indicated in the antibacterial activity assay were collected by centrifugation at 8000 rpm for 5 min and then resuspended in 0.9% NaCl. After washing with 0.9% NaCl three times, the bacteria cells were mixed with 2.5% (v/v) glutaraldehyde for 12 h to fix the cell morphology, followed by dehydration using graded ethanol (30%, 50%, 70%, 90%, and 95%). Then the bacterial cells were resuspended in 100% ethanol and dripped in a silicon slide for SEM imaging.

### Cytotoxicity Evaluation

The toxicity of sorbicillinoids towards AT-II (normal human lung cell) by MTT assay. AT-II cells were cultivated in Dulbecco's modified eagle medium (DMEM), containing 10% fetal bovine serum, 100 U/mL of penicillin, and 100 μg/mL streptomycin in an incubator with 5% CO_2_ at 37 °C. The cells were seeded in a 96-well plate at a density of 5000 cells per well. After 24 h of culture, the cells were mixed with sorbicillinoids dissolved in DMEM at the final concentrations of 0 50, 100, 150 200, 250, 300, 350, 400 and 450 μg/mL, followed by 24 h of culture. Then, 10 μL of MTT (5 mg/mL) was added into each well. After incubation for 4 h, the solution was removed and 150 μL DMSO was added into each cell. Then, the absorbance at 492 nm was measured with a microplate photometer (Multiskan FC, Thermo Fisher Scientific, USA).

### Hemolysis activity

Fresh blood was collected from a healthy male mouse and restored in an anticoagulation tube to avoid the blood coagulation. Red blood cells (RBCs) were obtained from the blood by centrifugation at 2000 rpm for 5 min and resuspended in 0.9% NaCl. The obtained RBCs were treated with different concentrations of sorbicillinoids (50, 100, 200, 250, and 300 μg/mL) for 2 h at 37 °C, and centrifuged at 2000 rpm for 5 min. Then the resultant supernatants were transferred to a 96-well plate for the absorbance measurement at 492 nm with a microplate photometer (Multiskan FC, Thermo Fisher Scientific, USA). RBCs treated with phosphate-buffered saline (PBS) and triton X-100 were set as the negative control and positive control, respectively. The hemolysis percentage was calculated by the following formula:

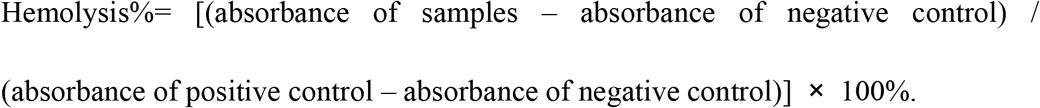

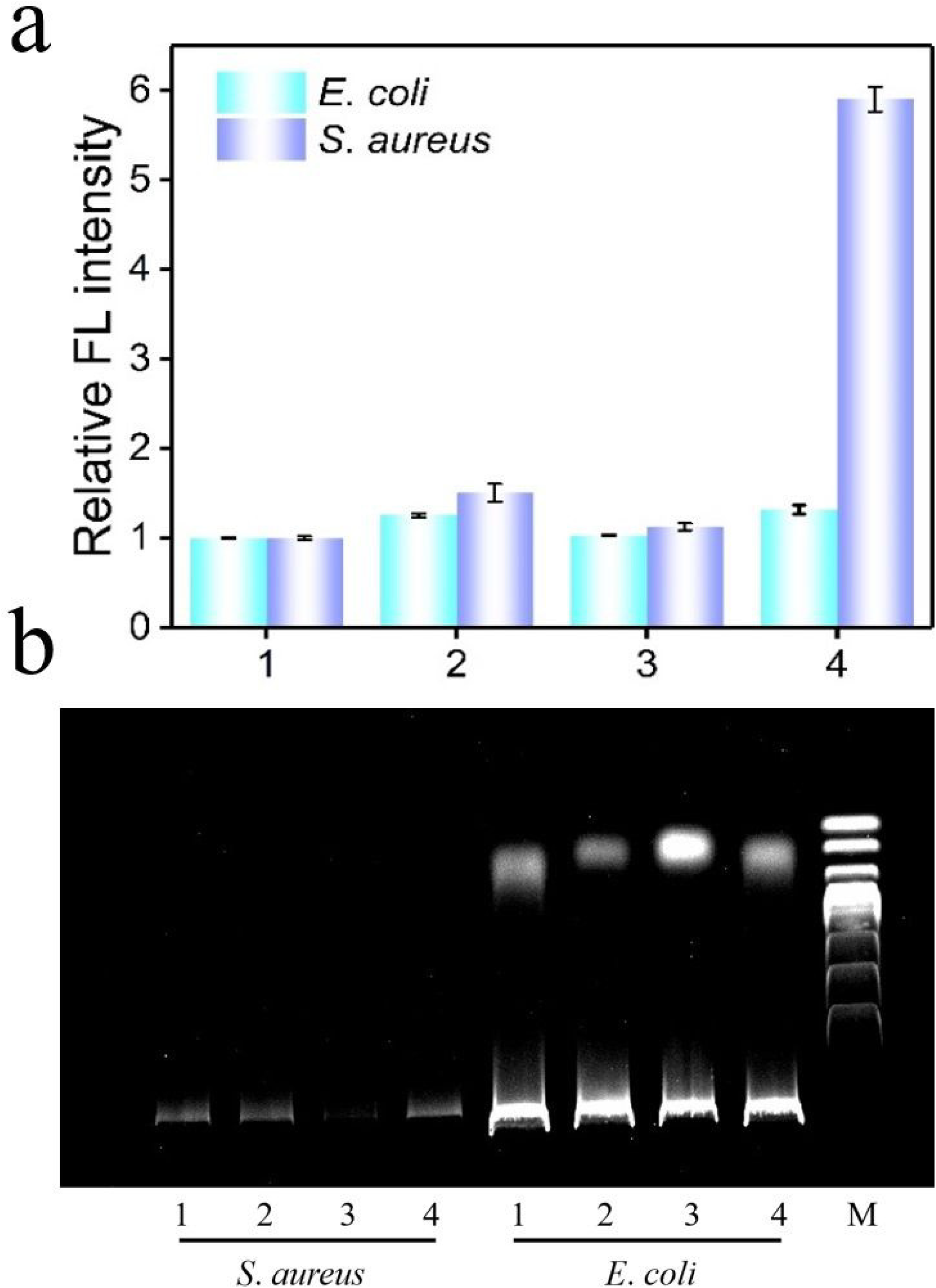

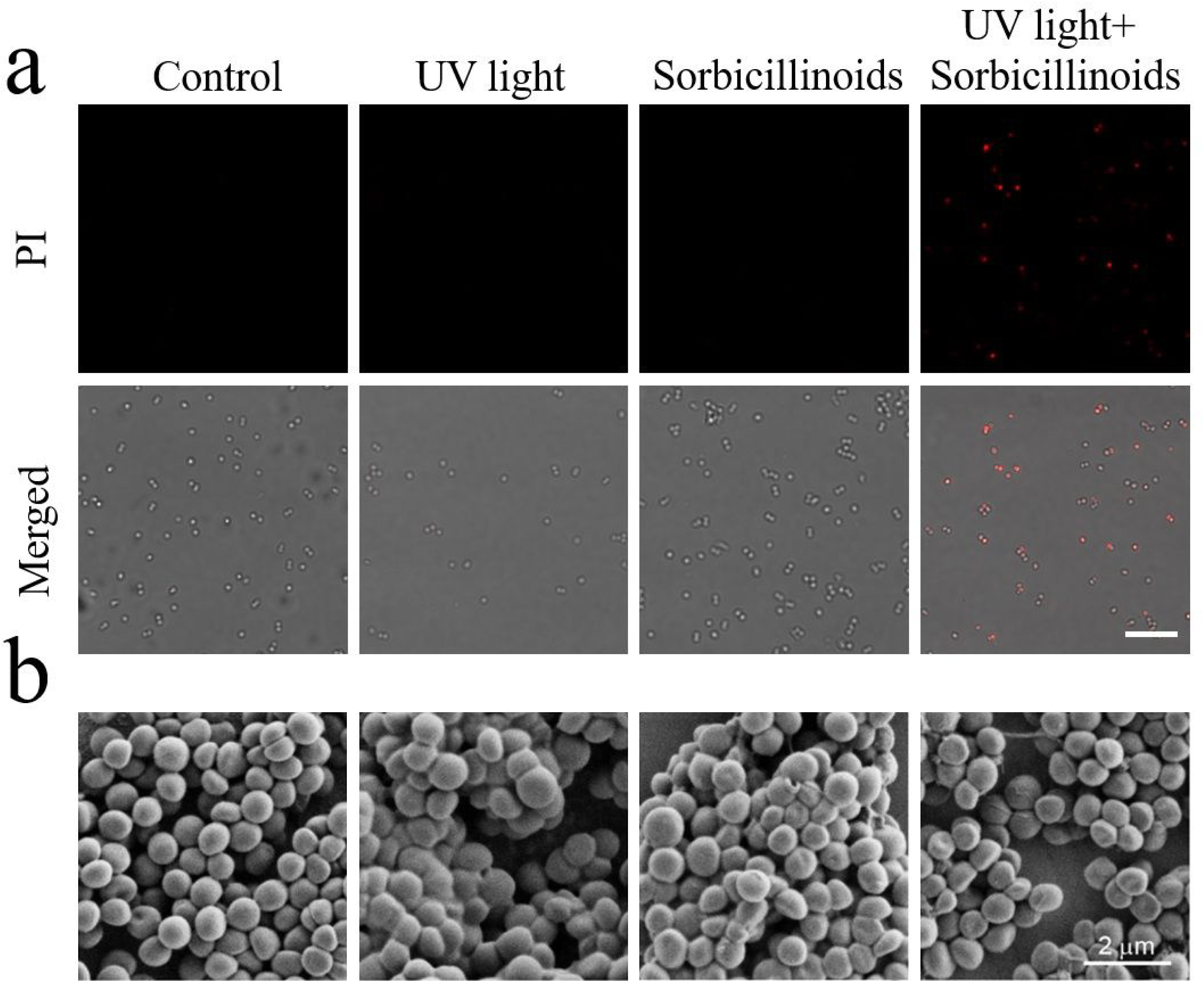

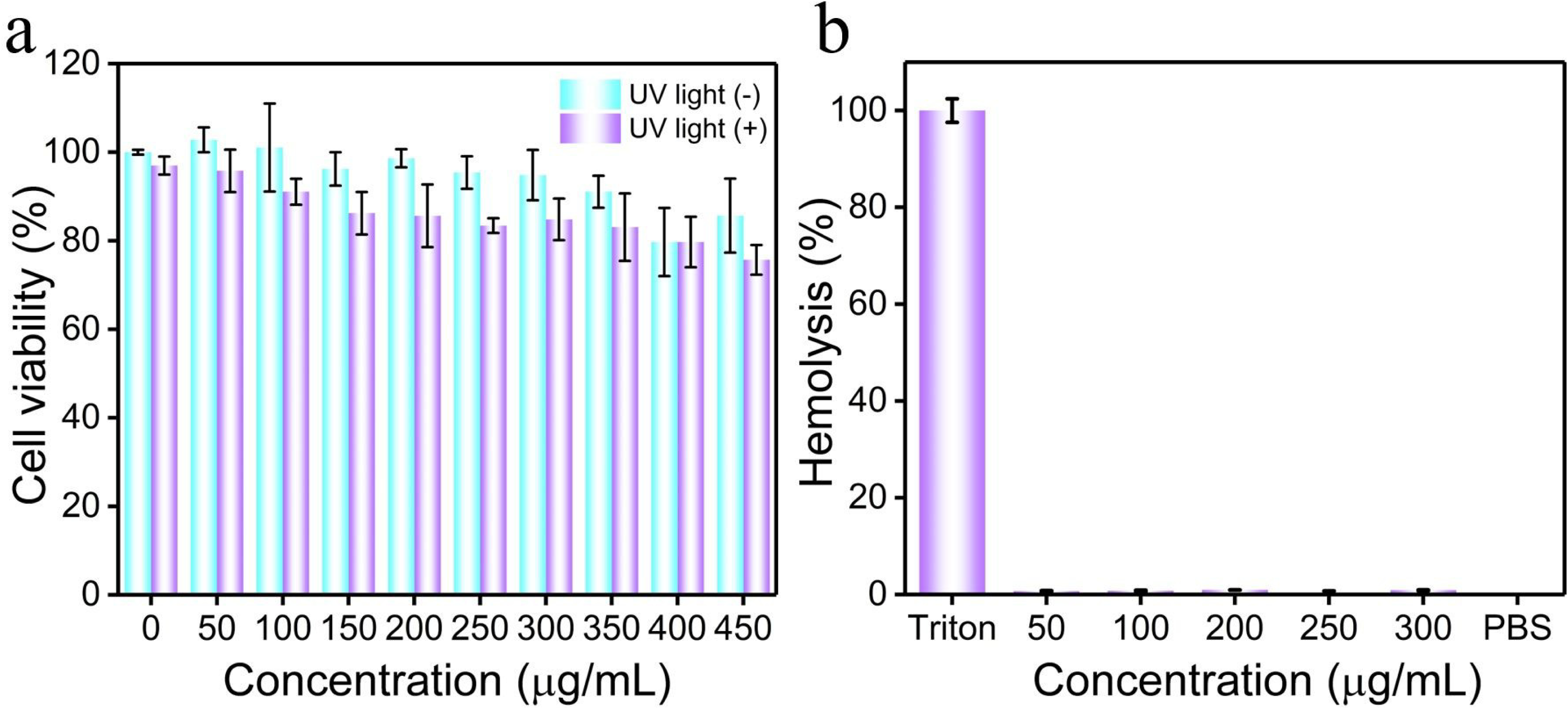

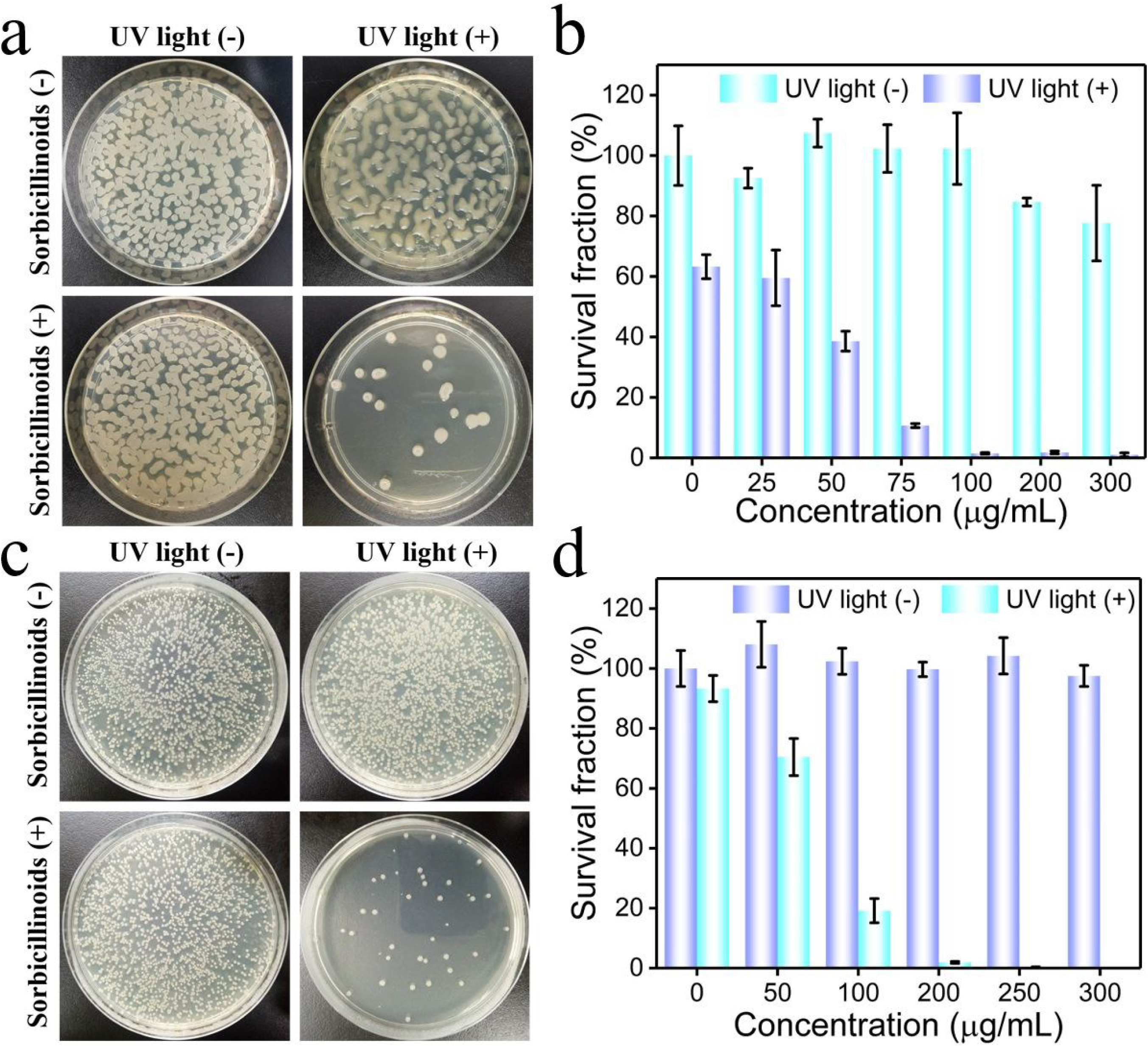

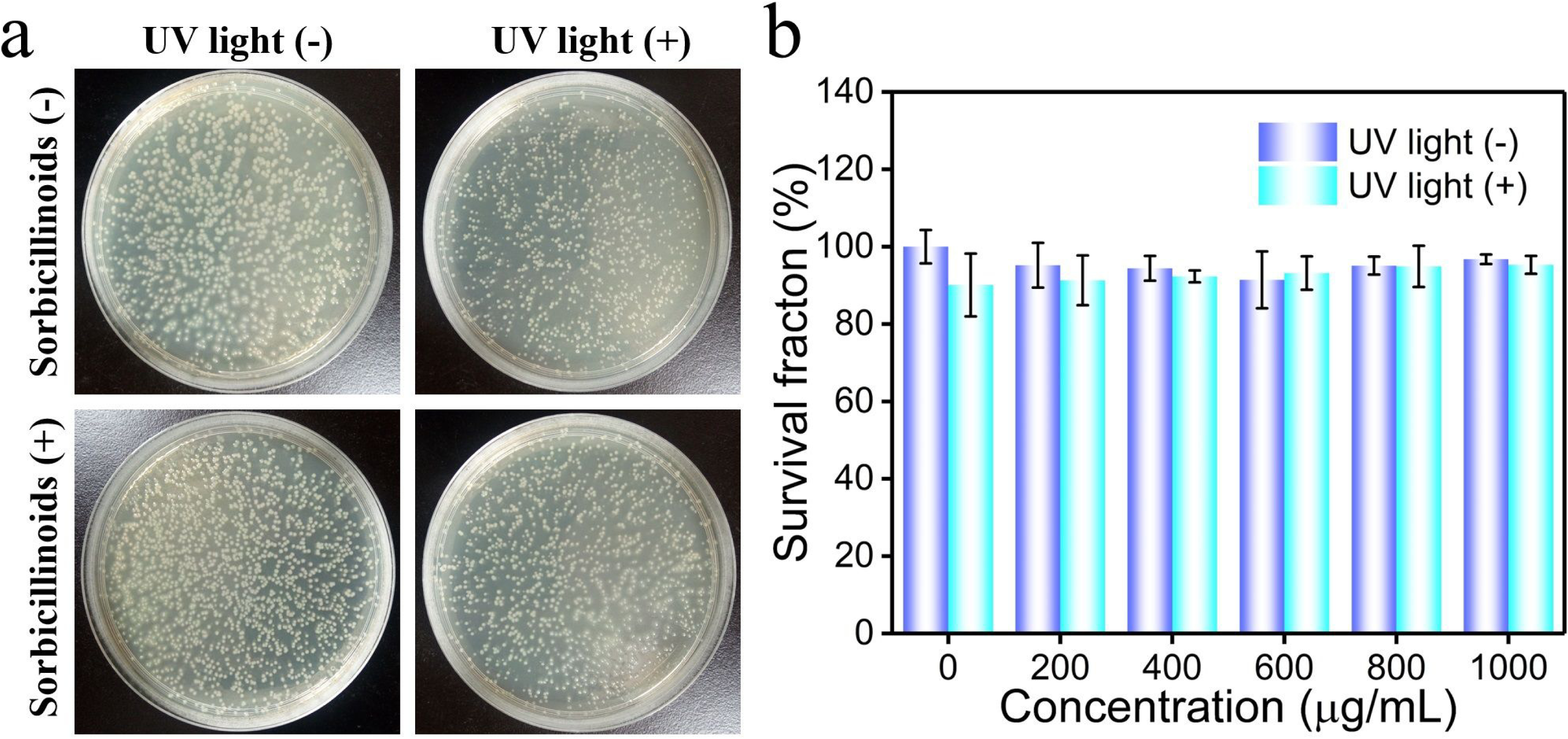

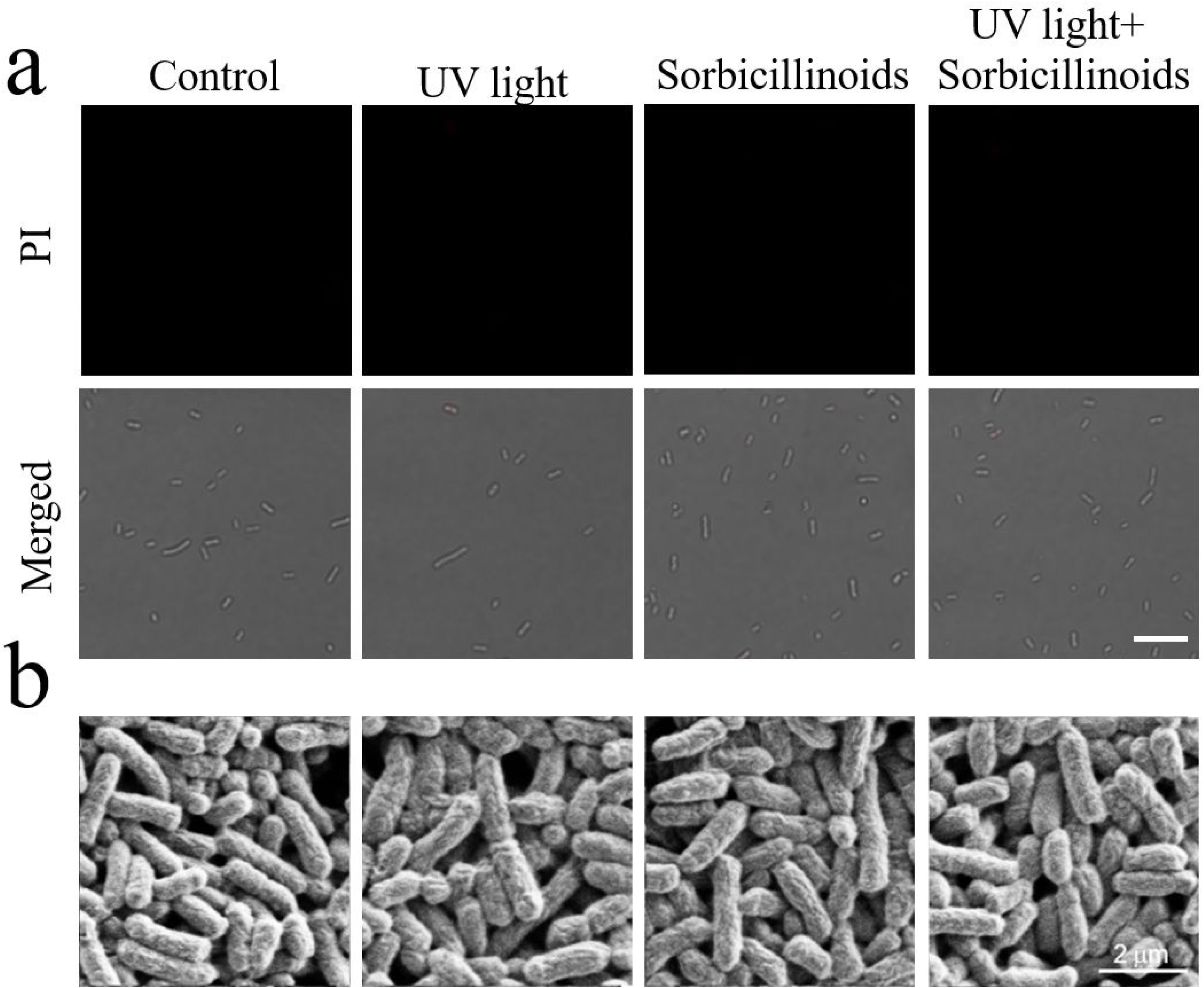

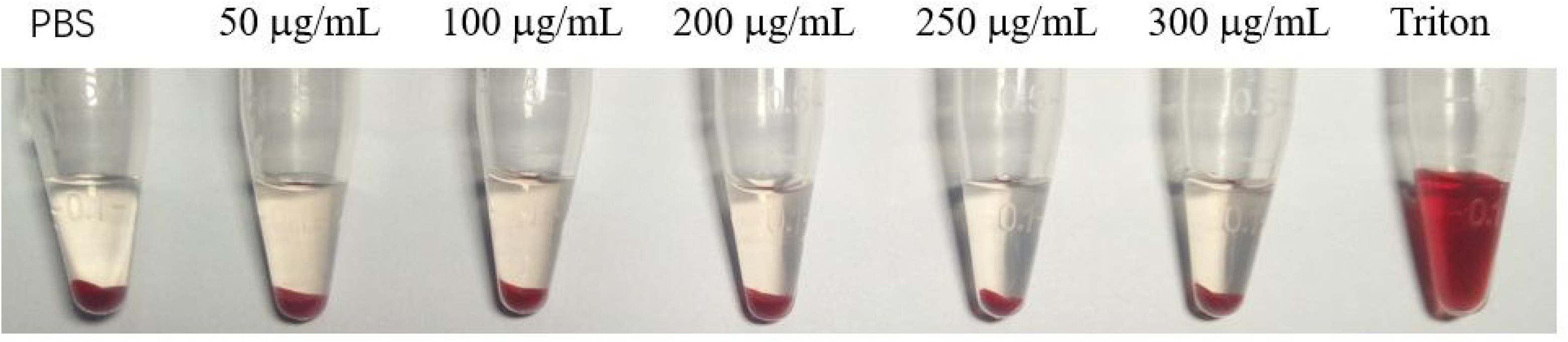

## Acknowledgements

This work was supported by grants from the National Natural Science Foundation of China (31700040), and the Fundamental Research Funds for the Central Universities.

## Authors’ contributions

YZHY and LFM designed this study; YZHY, LJY and QY performed the experiments; YZHY and LJY analyzed the data. YZHY wrote the manuscript. All authors read and approved the final manuscript.

## References

1. Jones KE, Patel NG, Levy MA, Storeygard A, Balk D, Gittleman JL, Daszak P. 2008. Global trends in emerging infectious diseases. Nature 451: 990–993.

2. Theuretzbacher U. 2013. Global Antibacterial resistance: the never-ending story. J. Glob Antimicrob Resist 1: 63–69.

3. Fischbach MA, Walsh CT. 2009. Antibiotics for emerging pathogens. Science 325: 1089–1093.

4. Taubes G. 2008. The bacteria fight back. Science 321: 356–361.

5. Levy SB, Marshall B. 2004. Antibacterial resistance worldwide: causes, challenges and responses. Nat Med 10: S122–S129.

6. Tenover FC. 2006. Mechanisms of antimicrobial resistance in bacteria. Am J Med 119: S3–S10.

7. Yang JJ, Zhang XD, Ma YH, Gao G, Chen XK, Jia HR, Li YH, Chen Z, Wu FG. 2016. Carbon dot-based platform for simultaneous bacterial distinguishment and antibacterial applications. ACS Appl Mater Interfaces 8: 32170–32181.

8. Ran HH, Cheng XT, Bao YW, Hua XW, Gao G, Zhang XD, Jiang YW, Zhu YX, Wu FG. 2019. Multifunctional quaternized carbon dots with enhanced biofilm penetration and eradication efficiencies. J Mater Chem B 7: 5104–5114.

9. Liu LH, Xu KJ, Wang HY, Tan JPK, Fan WM, Venkatraman SS, Li LJ, Yang YY. 2009. Self-assembled cationic peptide nanoparticles as an efficient antimicrobial agent. Nat Nanotechnol 4: 457–463.

10. Lam SJ, O’Brien-Simpson NM, Pantarat N, Sulistio A, Wong EHH, Chen YY, Lenzo JC, Holden JA, Blencowe A, Reynolds EC, Qiao GG. 2016. Combating multidrug-resistant Gram-negative bacteria with structurally nanoengineered antimicrobial peptide polymers. Nat Microbiol 1: 16162.

11. Hamblin MR, Hasan T. 2004. Photodynamic therapy: a new antimicrobial approach to infectious disease? Photochem Photobiol Sci 3: 436–450.

12. Liu Y, Qin R, Zaat SAJ, Breukink E, Heger M. 2015. Antibacterial photodynamic therapy: overview of a promising approach to fight antibiotic-resistant bacterial infections. J Clin Transl Res 1: 140–167.

13. Zhao Y, Dai XM, Wei XS, Yu YJ, Chen XL, Zhang XG, Li CX. 2018. Near-infrared light-activated thermosensitive liposomes as efficient agents for photothermal and antibiotic synergistic therapy of bacterial biofilm. ACS Appl Mater Interfaces 10: 14426–14437.

14. Wu SM, Li AH, Zhao XY, Zhang CL, Yu BR, Zhao NN, Xu FJ. 2019. Silica-coated gold-silver nanocages as photothermal antibacterial agents for combined anti-infective therapy. ACS Appl Mater Interfaces 11: 17177–17183.

15. Jia HR, Zhu YX, Chen Z, Wu FG. 2017. Cholesterol-assisted bacterial cell surface engineering for photodynamic inactivation of Gram-positive and Gram-negative bacteria. ACS Appl Mater Interfaces 9: 15943–15951.

16. Ravikumar M, Raghav D, Rathinasamy K, Kathiravan A, Mothi EM. 2018. DNA targeting long-chain alkoxy appended tin(IV) porphyrin scaffolds: photophysical and antimicrobial PDT investigations. ACS Appl Bio Mater 1: 1705–1716.

17. Zhang C, Gao F, Wu W, Qiu WX, Zhang L, Li RQ, Zhuang ZN, Yu WY, Cheng H, Zhang XZ. 2019. Enzyme-driven membrane-targeted chimeric peptide for enhanced tumor photodynamic immunotherapy. ACS Nano 13: 11249–11262.

18. Davies MJ. 2003. Singlet oxygen-mediated damage to proteins and its consequences. Biochem Biophys Res Commun 305: 761–770.

19. Alves E, Faustino MAF, Tomé JPC, Neves MGPMS, Tomé AC, Cavaleiro JAS, Cunha Â, Gomes NCM, Almeida A. 2013. Nucleic acid changes during photodynamic inactivation of bacteria by cationic porphyrins. Bioorg Med Chem 21: 4311–4318.

20. Almeida-Marrero V, van de Winckel E, Anaya-Plaza E, Torres T, de la Escosura A. 2018. Porphyrinoid biohybrid materials as an emerging toolbox for biomedical light management. Chem Soc Rev 47: 7369–7400.

21. Han K, Lei Q, Wang SB, Hu JJ, Qiu WX, Zhu JY, Yin WN, Luo X, Zhang XZ. 2015. Dual-stage-light-guided tumor inhibition by mitochondria-targeted photodynamic therapy. Adv Funct Mater 25: 2961–2971.

22. Parthasarathy A, Pappas HC, Hill EH, Huang Y, Whitten DG, Schanze KS. 2015. Conjugated polyelectrolytes with imidazolium solubilizing groups. properties and application to photodynamic inactivation of bacteria. ACS Appl Mater Interfaces 7: 28027–28034.

23. Malcher M, Volodkin D, Heurtault B, André P, Schaaf P, Möhwald H, Voegel JC, Sokolowski A, Ball V, Boulmedais F, Frisch B. 2008. Embedded silver ions-containing liposomes in polyelectrolyte multilayers: cargos films for antibacterial agents. Langmuir 24: 10209–10215.

24. Huh AJ, Kwon YJ. 2011. “Nanoantibiotics”: A new paradigm for treating infectious diseases using nanomaterials in the antibiotics resistant era. J Control Release 156: 128–145.

25. Wegener M, Hansen MJ, Driessen AJM, Szymanski W, Feringa BL. 2017. Photocontrol of antibacterial activity: shifting from UV to red light activation. J Am Chem Soc 139: 17979–17986.

26. García I, Ballesta S, Gilaberte Y, Rezusta A, Pascual Á. 2015. Antimicrobial photodynamic activity of hypericin against methicillin-susceptible and resistant *Staphylococcus aureus* biofilms. Future Microbiol 10: 347–356.

27. Tonon CC, Paschoal MA, Correia M, Spolidório DMP, Bagnato VS, Giusti JSM, Santos-Pinto L. 2015. Comparative effects of photodynamic therapy mediated by curcumin on standard and clinical isolate of *Streptococcus mutans*. J Contemp Dent Pract 16: 1–6.

28. Zhenjun D, Lown JW. 1990. Hypocrellins and their use in photosensitization. Photochem Photobiol 52: 609–616.

29. Maisch T, Eichner A, Späth A, Gollmer A, König B, Regensburger J, Bäumler W. 2014. Fast and effective photodynamic inactivation of multiresistant bacteria by cationic riboflavin derivatives. PLoS One 9: e111792.

30. Abrahamse H, Hamblin MR. 2016. New photosensitizers for photodynamic therapy. Biochem J 473: 347–364.

31. Meng JJ, Wang XH, Xu D, Fu XX, Zhang XP, Lai DW, Zhou LG, Zhang GZ. 2016. Sorbicillinoids from fungi and their bioactivities. Molecules 21: 715

32. Harned AM, Volp KA. 2011. The sorbicillinoid family of natural products: isolation, biosynthesis, and synthetic studies. Nat Prod Rep 28: 1790–1810.

33. Abe NHirota, A. 2002. Chemical studies of the radical scavenging mechanism of bisorbicillinol using the 1,1-diphenyl-2-picrylhydrazyl radical. Chem Commun 6: 662–663.

34. Kawahara T, Takagi M, Shin-ya K. 2012. JBIR-124: A Novel Antioxidative agent from a marine sponge-derived fungus *Penicillium citrinum* SpI080624G1f01. J Antibiot 65: 45–47.

35. Guo WQ, Zhang ZZ, Zhu TJ, Gu QQ, Li DH. 2015. Penicyclones A-E, antibacterial polyketides from the deep-sea-derived fungus *Penicillium sp*. F23-2. J Nat Prod 78: 2699–2703.

36. Meng JJ, Cheng W, Heydari H, Wang B, Zhu K, Konuklugil B, Lin WH. 2018. Sorbicillinoid-based metabolites from a sponge-derived fungus *Trichoderma saturnisporum*. Marine Drugs 16: 226

37. Lai DW, Meng JJ, Zhang XP, Xu D, Dai JG, Zhou LG. 2019. Ustilobisorbicillinol A, a cytotoxic sorbyl-containing aromatic polyketide from *Ustilaginoidea virens*. Org Lett 21: 1311–1314.

38. Meng JJ, Gu G, Dang PQ, Zhang XP, Wang WX, Dai JG, Liu Y, Lai DW, Zhou LG. 2019. Sorbicillinoids from the fungus *Ustilaginoidea virens* and their phytotoxic, cytotoxic, and antimicrobial activities. Frontiers in Chemistry 7: 435

39. Minty JJ, Singer ME, Scholz SA, Bae CH, Ahn JH, Foster CE, Liao JC, Lin XN. 2013. Design and characterization of synthetic fungal-bacterial consortia for direct production of isobutanol from cellulosic biomass. Proc Natl Acad Sci 110: 14592–14597.

40. Li CC, Lin FM, Sun W, Yuan SX, Zhou ZH, Wu FG, Chen Z. 2018. Constitutive hyperproduction of sorbicillinoids in *Trichoderma reesei* ZC121. Biotechnol Biofuels 11: 291.

41. Derntl C, Kluger B, Bueschl C, Schuhmacher R, Mach RL, Mach-Aigner AR. 2017. Transcription factor Xpp1 is a switch between primary and secondary fungal metabolism. Proc Natl Acad Sci 114: 560–569.

42. Mao CY, Xiang YM, Liu XM, Cui ZD, Yang XJ, Yeung KWK, Pan HB, Wang XB, Chu PK, Wu SL. 2017. Photo-inspired antibacterial activity and wound healing acceleration by hydrogel embedded with Ag/Ag@AgCl/ZnO nanostructures. ACS Nano 11: 9010–9021.

43. Ran CZ, Zhang ZD, Hooker J, Moore A. 2012. In Vivo Photoactivation without “light”: use of cherenkov radiation to overcome the penetration limit of light. Mol Imaging Biol 14: 156–162.

44. Lien EJC, Hansch C, Anderson SM. 1968. Structure-activity correlations for antibacterial agents on Gram-positive and Gram-negative cells. J Med Chem 11: 430–441.

45. Moan J, Berg K. 1991. The photodegradation of porphyrins in cells can be used to estimate the lifetime of singlet oxygen. Photochem Photobiol 53: 549–553.

46. Cieplik F, Deng DM, Crielaard W, Buchalla W, Hellwig E, Al-Ahmad A, Maisch T. 2018. Antimicrobial photodynamic therapy - what we know and what we don’t. Crit Rev Microbiol 44: 571–589.

47. Mao CY, Xiang YM, Liu XM, Zheng YF, Yeung KWK, Cui ZD, Yang XJ, Li ZY, Liang YQ, Zhu SL, Wu SL. 2019. Local photothermal/photodynamic synergistic therapy by disrupting bacterial membrane to accelerate reactive oxygen species permeation and protein leakage. ACS Appl Mater Interfaces 11: 17902–17914.

48. Li CC, Lin FM, Sun W, Wu FG, Yang H, Lv RJ, Zhu YX, Jia HR, Wang C, Gao G, Chen Z. 2018. Self-assembled rose bengal-exopolysaccharide nanoparticles for improved photodynamic inactivation of bacteria by enhancing singlet oxygen generation directly in the solution. ACS Appl Mater Interfaces 10: 16715–16722.

